# The dual role of a highly structured RNA (the S fragment) in the replication of foot-and-mouth disease virus

**DOI:** 10.1101/2023.04.26.538422

**Authors:** Joseph C. Ward, Lidia Lasecka-Dykes, Samuel J. Dobson, Sarah Gold, Natalie J. Kingston, Morgan R. Herod, Donald P. King, Tobias J. Tuthill, David J. Rowlands, Nicola J. Stonehouse

## Abstract

Secondary and tertiary RNA structures play key roles in genome replication of single stranded positive sense RNA viruses. Complex, functional structures are particularly abundant in the untranslated regions of picornaviruses, where they are involved in initiation of translation, priming of new strand synthesis and genome circularisation. The 5′ UTR of foot-and-mouth disease virus (FMDV) is predicted to include a *c.* 360 nucleotide-long stem-loop, termed the short (S) fragment. This structure is highly conserved and essential for viral replication, but the precise function(s) are unclear. Here, we used selective 2′ hydroxyl acetylation analysed by primer extension (SHAPE) to experimentally-determine aspects of the structure, alongside comparative genomic analyses to confirm structure conservation from a wide range of field isolates. To examine its role in virus replication, we introduced a series of deletions to the distal and proximal regions of the stem loop. These truncations affected genome replication in a size-dependent and, in some cases, host cell-dependent manner. Furthermore, during passage of viruses incorporating the largest tolerated deletion from the proximal region of the S fragment stem loop, an additional mutation was selected in the viral RNA-dependent RNA polymerase, 3D^pol^.These data suggest that the S fragment and 3D^pol^ interact in the formation of the FMDV replication complex.

## Introduction

The genomes of single-stranded positive sense RNA viruses include both coding and regulatory regions. Picornaviruses are amongst the smallest, with genomes approximately 7.5 - 8 kb in length. Typically, the genome comprises a major single open reading frame flanked by 5′ and 3′ untranslated regions (UTRs), however, additional small open reading frames have been identified in some cardioviruses and enteroviruses (1, 2). The large 5′ UTRs present in most picornaviruses range from ∼800 to ∼1300 nucleotides and have been predicted *in silico* by energy minimisation algorithms to comprise several distinct highly structured domains. Some of these RNA structures are common for all picornaviruses and have been well characterised, for example an internal ribosome entry site (IRES) responsible for the initiation of translation, which comprises ∼450 nucleotides for most picornaviruses (although smaller IRES elements resembling those found in some flaviviruses were also found) (3–5). The number and organisation of these structural domains varies amongst picornaviruses. For example, a small stem-loop, the cis-active replicative element (*cre*), which is essential for replication, is found in the 5′ UTR of aphthoviruses, but within the coding region in enteroviruses (6–8). All picornavirus genomes include a structured domain at the 5′ end, although the size, conformation and sequence varies in viruses from different genera. Enterovirus 5′ UTRs terminate in a ∼80 nucleotide cloverleaf structure, while in some other picornaviruses the 5′ end forms an extended stem-loop (9–11). The size of this stem-loop varies markedly; 40 nucleotides in hepatoviruses and ∼80 nucleotides in cardioviruses, whereas the largest (∼360 nucleotides) is present in the aphthoviruses (12). The functions of some of these RNA structures remain unknown.

Here, we focus on the long 5′ terminal stem-loop (termed S fragment) of foot-and-mouth disease virus (FMDV), an aphthovirus which infects cloven-hoofed animals. It is the causative agent of foot-and-mouth disease (FMD), a highly contagious and economically important infection posing a constant threat to the global livestock industry. In endemic countries, FMD is controlled by vaccination and by movement restrictions, while movement restrictions and slaughter have been used when outbreaks occurred in non-endemic countries (13). Control by vaccination is complicated by high antigenic variability and the occurrence of asymptomatic carrier animals (14–16). FMDV has an 8.5 kb RNA genome organised into L, P1-2A, P2 and P3 coding regions, flanked by the 5′ UTR described above and a short 3′ UTR. L is a protease responsible for separating itself from the polyprotein and cleaving key cellular proteins, P1 encodes the structural (capsid) proteins, while P2 and P3 encode non-structural proteins involved in genome replication. The latter include the RNA-dependent RNA polymerase (RdRp, also termed 3D^pol^) and the protease 3C^pro^ alongside less well-characterised proteins including 2C, a potential helicase (17–19). At approximately 1.3 kb, the 5′ UTR of FMDV is ∼1/7^th^ of the entire genome and is the longest 5′ UTR within the picornavirus family (10). Its sequence comprises at least five structurally and functionally distinct domains. The S fragment stem-loop is located at the 5′ terminus and is followed by a long poly-C-tract of variable length (about 70 to 250 nucleotides), 2-4 tandem repeated sequences predicted to form pseudoknots, the small stem-loop *cre* involved in uridylation of the replication primer peptide VPg and finally the IRES element responsible for protein translation initiation (20). The functions of several of these domains are poorly understood (21, 22), although we have recently described the importance of a sequence in the pseudoknot region for virus assembly (23).

Whilst the S fragment is known to be essential for replication, its precise function remains unclear. The secondary structure of the S fragment appears to be conserved, although its sequence and length varies between different clades of FMDV (10, 24). Truncated S fragments have been observed for some field isolates, but these retain the long stem-loop structure (10). It has been reported that while distal portions of this stem-loop structure are not essential for genome replication, they play a role in the modulation of host-cell innate immune responses (25, 26). The 5’ end of poliovirus (PV) RNA also terminates in a highly structured region, called the cloverleaf, which is involved in the switch from protein translation to RNA replication, in addition to recruiting host and viral proteins which contribute to RNA circularisation during genome replication (9, 27). It is possible that the S fragment in FMDV has similar functions; it is known to interact with the 3′ UTR, suggesting a role in genome circularisation (28).

We used selective-2′ hydroxyl acylation analysed by primer extension (SHAPE) to provide direct evidence to complement comparative bioinformatic analyses and *in silico* predictions of the structure of the S fragment. We confirm that substantial deletions can be made to the distal region of the S fragment stem-loop without seriously compromising *in vitro* replication, although host cell differences were observed. We also show that proximal deletions are less well tolerated, which correlates with our observation that the proximal part of the S fragment shows higher conservation of nucleotide pairings. Furthermore, viruses reconstructed to include the maximal viable proximal deletion can, during sequential passages, select a compensatory mutation in the 3D^pol^. This mutation is located in a highly conserved position known to interact with template RNA, and enhanced replication of a mutant replicon carrying this proximal S fragment deletion.

## Materials and Methods

### Cells lines

Baby hamster kidney (BHK)-21 and Madin-Darby bovine kidney (MDBK) cells were obtained from the American Type Culture Collection (ATCC) (LGC Standard) and maintained in Dulbecco’s modified Eagle’s Medium with glutamine (Sigma-Aldrich) supplemented with 10% foetal calf serum (FCS), 50 U/ml penicillin and 50 µg/ml streptomycin as previously described (29).

### Plasmid construction

The FMDV ptGFP replicon plasmids along with the equivalent 3D^pol^ knock-out controls (3D^pol^ -GNN) have already been described (30, 31).

A variant of the ptGFP replicon was produced to allow for easy mutagenesis of the S-fragment. An *Eag*I site was introduced at each end of the S-fragment by PCR mutagenesis, altering the wild-type sequence from AAAGGGGGCATTA to AAACGGCCGATTA at the 5′ end and GCGCCCGCCTTT to GCGCGGCCGTTT at the 3′ end. S-fragment sequences containing deletions were chemically synthesised (Thermo Fisher Scientific) and inserted into the replicon vector using complementary *Eag*I sites.

Mutation of residue I189L of 3D^pol^ in the ptGFP replicon was achieved by PCR mutagenesis resulting in A7023C substitution. All primer sequences are available on request.

### Protein purification

The I189L mutation was introduced into the His-tagged 3D^pol^ expression clone pET28a-3D, and recombinant protein was expressed and purified as previously described (32, 33).

### *In vitro* transcription

*In vitro* transcription reactions were performed as described previously (34, 35). Briefly, 5 µg of replicon plasmid was linearised with *Asc*I (NEB), purified by phenol-chloroform extraction, ethanol precipitated, redissolved in RNase-free water and used in a T7 *in vitro* transcription reaction. Reactions were incubated at 32°C for 4 hours, treated with 1.25 units of RQ1 DNase for 30 minutes at 37°C and RNA recovered using an RNA Clean & Concentrator-25 spin column kit (Zymo Research), following manufacturer’s instructions. All transcripts were quantified by NanoDrop 1000 (Thermo Scientific) and RNA integrity assessed by 3-(N-morpholino) propanesulfonic acid (MOPS) -formaldehyde gel electrophoresis.

### Replication assays

BHK-21 and MDBK cell based replicon replication assays were performed in 24-well plates with 0.5 µg/cm^2^ of RNA using Lipofectin transfection reagent (Life Technologies) as previously described (35). For every experiment, transfection was performed in duplicate and experiments biologically repeated as indicated. Replicon replication was assessed by live cell imaging using an IncuCyte Zoom Dual colour FLR, an automated phase-contrast and fluorescent microscope within a humidifying incubator. At hourly intervals up to a defined end point, images of each well were taken and used to enumerate the ptGFP positive cell count per well, using our established protocols (34, 35). Data are shown at 8 hours post transfection (when maximum replication was observed) on a linear scale.

### SHAPE analysis

RNA transcripts representing the entire 5′ UTR of FMDV (nucleotides 1 – 1581) were prepared as above. A sample containing 12 pmol of transcribed RNA was heated to 95°C for 2 minutes before cooling on ice. RNA folding buffer (100 mM HEPES, 66 mM MgCl_2_ and 100 mM NaCl) was added to the RNA and incubated at 37°C for 30 minutes. The folded RNA was treated with 5 mM *N*-methyl isatoic anhydride (NMIA) or dimethyl sulfoxide (DMSO) for 50 minutes at 37°C. The chemically modified RNA was ethanol precipitated and resuspended in 0.5x Tris-EDTA (TE) buffer.

Hex or FAM fluorescent primers were bound to modified RNA by heating the reaction to 85°C for 1 minute, 60°C for 10 minutes and 35°C for 10 minutes in a thermocycler. Reverse transcription was continued using Superscript III (Invitrogen) following manufacturers protocol with incubation at 52°C for 30 minutes.

Post-extension, cDNA:RNA hybrids were disassociated by incubation with 4M NaOH at 95°C for 3 minutes before neutralisation with 2M HCl. Extended cDNA was ethanol precipitated and resuspended in deionized formamide (Thermo Fisher Scientific). Sequencing ladders were similarly produced using 6 pmol of RNA with the inclusion of 10 mM ddCTP in the reverse transcription mix and using a differentially labelled fluorescent primer (either Hex or FAM). A sequencing ladder was combined with NMIA or DMSO samples and dispatched on dry ice for capillary electrophoresis (DNA Sequencing and Services, University of Dundee) (23).

NMIA reactivity was used as constraints for RNA secondary structure prediction of the S fragment using the Vienna RNA probing package and based on sequence of the replicon used in this study (36).

### Construction of recombinant viruses

Replicons used here are based on plasmid pT7S3 which encodes a full-length infectious copy of FMDV O1 Kaufbeuren (O1K) (37). To generate infectious viral genomes the reporter sequence was removed from replicons by digestion with flanking *Psi*I and *Xma*I restriction enzymes and replaced with the corresponding fragment from pT7S3 encoding the capsid proteins. Full length viral RNA was transcribed using a T7 MEGAscript kit (Thermo Fisher Scientific), DNase treated using TurboDNase (Thermo Fisher Scientific) and purified using a MEGAclear Transcription Clean-Up kit (Thermo Fisher Scientific). RNA quality and concentration were determined by MOPS-formaldehyde denaturing agarose gel electrophoresis and Qubit RNA BR Assay Kit (Thermo Fisher Scientific).

### Virus recovery

BHK-21 cells were transfected in 25 cm^2^ flasks with 8 µg per flask of infectious clone-derived RNA using TransIT transfection reagent (Mirus) as described previously (35). At full cytopathic effect (CPE) or 24 hours post-transfection (whichever was earlier) cell lysates were freeze-thawed and clarified by centrifugation. The clarified lysate (1 ml) was blind passaged onto naïve BHK-21 cells and this was repeated for five rounds of passaging.

### Sequencing of recovered virus

Recovered viruses at passage 3, 4 and 5, were sequenced with an Illumina Miseq (Illumina) using a modified version of a previously described PCR-free protocol (38). Total RNA was extracted from clarified lysates using TRizol reagent (Thermo Fisher Scientific) and residual genomic DNA was removed using DNA-free DNA removal Kit (Thermo Fisher Scientific) prior to ethanol precipitation. Purified RNA was used in a reverse transcription reaction as previously described (38, 39). Following reverse transcription, cDNA was purified and quantified using a Qubit ds-DNA HS Assay kit (Thermo Fisher Scientific) and a cDNA library prepared using Nextera XT DNA Sample Preparation Kit (Illumina). Sequencing was carried out on the MiSeq platform using MiSeq Reagent Kit v2 (300 cycles) chemistry (Illumina) and paired-end sequencing.

FastQ files were quality checked using FastQC with poor quality reads filtered using the Sickle algorithm (40). Host cell reads were removed using FastQ Screen algorithm and FMDV reads assembled *de novo* into contigs using IDBA-UD (41). Contigs that matched the FMDV library (identified using Basic Local Alignment Search Tool (BLAST)) were assembled into consensus sequences using SeqMan Pro software in the DNA STAR Lasergene 13 package (DNA STAR) (42). Finally, the filtered fastQ reads were aligned to the *de novo* constructed consensus sequences using BWA-MEM algorithm incorporated into Burrows-Wheeler Aligner (BWA), and mutations were visualised using the Integrative Genomics Viewer (IGV) (43–46).

### Cell killing assays

Virus titre was determined by plaque assay on BHK-21 cells as described before (47). BHK-21 cells were seeded with 3 x10^4^ cells/well in 96 well plates and allowed to settle overnight. Cell monolayers were inoculated with each rescued virus at MOI of 0.01 PFU for 1 hour, inoculum was removed and 150 µl of fresh GMEM (supplemented with 1% FCS) was added to each well. Appearance of CPE was monitored every hour by live cell microscopy (contrast phase) using the IncuCyte S3. CPE was observed as rounding of the cells and progress of infection was monitored as a drop in confluency compared to the mock treated cells (i.e., treatment with uninfected cell lysate). Serial images of cells were analysed using IncuCyte S3 2018B software.

### Sym/Sub polymerase activity assays

Sym/Sub RNA with sequence 5 [GCAUGGGCCC 3 [was synthesised (Sigma Aldrich) and 5′ end labelled using γ^32^P UTP (Perking Elmer) and T4 polynucleotide kinase (NEB) following manufacturer’s protocol. Labelled RNA was purified by ethanol precipitation and resuspended in nuclease free water.

0.5 μM of end labelled RNA was incubated in polymerisation buffer (30 mM MOPS pH 7.0, 33 mM NaCl, 5 mM MgAc) and heated to 95°C for 2 minutes before chilling on ice. 2 μM of recombinant 3D^pol^ was added to the RNA and incubated at 37°C for 10 minutes to promote annealing. After annealing, 50 μM of rNTP was added to the reaction, addition of nucleotide varied depending on experiment. Aliquots were taken periodically, and reactions stopped by addition of 2x TBE-urea RNA loading dye (Thermo Fisher Scientific).

Extension products in loading dye were heated to 70°C for 5 minutes before loading on a 23% denaturing polyacrylamide gel containing 7 M urea. Samples were electrophoresed until sufficiently separated. After electrophoresis gels were fixed for 30 minutes in fixative solution and exposed onto a phosphorscreen (48, 49).

### Bioinformatic analysis

Full genome sequences of 118 FMDV isolates, representing all seven serotypes (Supplementary Table S1), were downloaded from GenBank. The sequences were chosen (based on the region encoding the VP1 protein) to represent the known genomic diversity of FMDV across all seven serotypes. VP1 is the most variable part of the FMDV genome and frequently used to calculate phylogenetic relationship of FMDV isolates.

The RNA structure of the S fragment was predicted as described before using the RNAalifold program implemented in the ViennaRNA package (36, 50). For the covariance analysis, also calculated using RNAalifold program, only those isolates were included which contained complete sequence at the 5′ end of the genome. Data representing covariance were superimposed onto the schematics of the S fragment RNA structure and visualised using Forna tool implemented in the ViennaRNA Web Services (51). The sequence logo representing amino acid conservation within positions 185 – 194 of the 3D^pol^ was prepared using a WebLogo 3.7.4 server and using 1123 FMDV 3D^pol^ sequences obtained from fmdbase.org (an FMDV sequence database generated by the FAO World Reference Laboratory for FMD at The Pirbright Institute) (52, 53).

## Results

### SHAPE analysis of the S fragment

Based on computational folding, such as mFOLD algorithms, the S fragment is predicted to form a single large hairpin stem-loop comprising approximately 360 nucleotides. Here, we use selective 2′ hydroxyl acylation analysed by primer extension (SHAPE) to further investigate the structure of this part of the FMDV genome. This approach relies on the formation of 2′-O-adducts in single stranded and accessible regions of the RNA, which can then be detected by reverse transcription.

RNA transcripts representing the full FMDV 5′ UTR were folded before incubation with 5 mM NMIA. The modified RNAs were then used in reverse-transcription reactions containing fluorescently end-labelled primers. Reverse transcription is terminated when the enzyme reaches a nucleotide with a NMIA adduct, thus creating a series of cDNA fragments of different lengths. These cDNA fragments were analysed by capillary gel electrophoresis alongside a sequencing ladder to identify the sites of termination, indicating the locations of unpaired and accessible nucleotides. The proximity of the poly-C-tract to the 3′ end of the S fragment limited potential primer binding sites, resulting in unreliable data for the last 60 nucleotides of the S fragment which were excluded from the analysis (shown in grey in Figure 1A). The rest of the S fragment was well covered and the SHAPE reactivities from 6 independent experiments were used to complement the mFOLD algorithm analyses to generate a structural prediction of this region (Figure 1).

**Figure 1.**
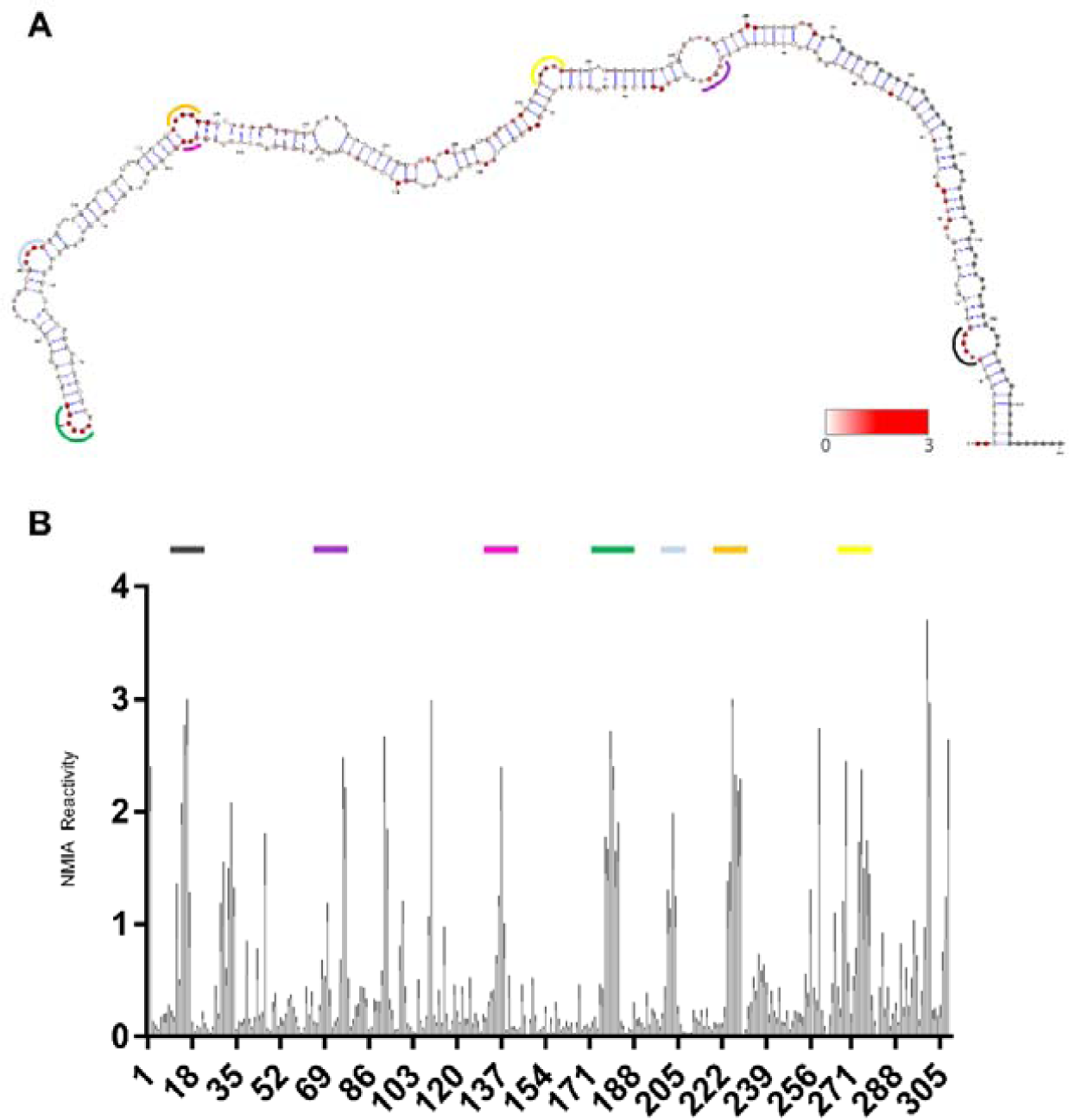
SHAPE analysis of the S fragment. **(A)** Superimposition of SHAPE reactivity on the *in silico* predicted secondary structure WT S fragment. Secondary structure was predicted using the Vienna RNA probing package and visualised using VARNA. NMIA reactivity is overlaid and represented on a colour scale from low (white) to high (red). Nucleotides for which there is no data are represented as grey. Coloured regions corresponding to the peaks in NIMIA reactivity are shown for ease of interpretation and localisation. **(B)** Individual nucleotide NMIA reactivity as analysed by SHAPE reactions and capillary electrophoresis. High reactivity indicates high probability of single-stranded regions i.e., non base-paired nucleotides. Data was analysed using QuSHAPE. Corresponding coloured regions are shown for some regions across **A** and **B** for ease of interpretation and localisation of SHAPE data to RNA structure. (n = 6, error bars represent SEM).

In general, the SHAPE mapping data fitted well with the *in silico* predictions with the most reactive residues coinciding with bulges within the predominately double stranded stem-loop structure (Figure 1A). The locations of reactive residues can be seen in the NMIA reactivity graph, where groups of nucleotides within bulges showed high reactivity (Figure 1B). However, some predicted bulges showed little NMIA reactivity, suggesting steric hindrance possibly due to higher order tertiary structure. Similarly, some predicted base-paired nucleotides were NMIA reactive. These mostly occurred at the tops and bottom of bulges and may result from nucleotides becoming transiently available for modification during ‘breathing’ (54). The 5′ terminal two nucleotides (uridines at position 1 and 2) were highly reactive and therefore likely to be non-base paired.

### The effects of deletions to the distal or proximal regions of the S fragment stem-loop on replicon replication

The SHAPE data and the structural prediction described in Figure 1 were used to design a series of truncations to the S fragment, and the modified sequences were introduced into a replicon. The replicon was based on an infectious clone of FMDV O1K in which the P1 region of the genome is replaced with a ptGFP reporter (55) (Figure 2A). Deletions were designed to end at either the top or bottom of bulges, helping ensure that the stem-loop hairpin structure was maintained. To facilitate replacement of the S fragment with modified sequences *Eag*I restriction sites were introduced at each end of the S fragment sequence. These sites required minimal modification of the original sequence and showed no predicted change in RNA structure, and the modified genome sequence is hereafter referred to as WT. Addition of the *Eag*I site at the 5′ end of the S fragment resulted in a large reduction in replication, however, when paired with the 3′ *Eag*I site, restoring base pairing, replicative potential was drastically improved, although still below that of the original O1K WT sequence (Figure S1). Deletions of 70, 148, 246 and 295 nucleotides were introduced at the distal end (top) of the S fragment stem-loop (constructs named D-70, D-148, D-246 and D-295, respectively) and deletions of 48, 97, 195 and 271 nucleotides were made to the proximal end (bottom) of the S fragment (constructs referred to as P-48, P-97, P-195 and P-271, respectively) (Figure 2B). The truncated sequences were chemically synthesised and introduced into the WT replicon via the *Eag*I sites, meaning that the highly conserved 5′ and 3′ regions and essential UU nucleotides at positions 1 and 2 were maintained in all truncation mutants. The eight new replicon clones were transcribed into RNAs and transfected into BHK-21 or MDBK cells, alongside WT and a 3D^pol^-GNN negative control. The 3D^pol^-GNN replicon cannot replicate and so the ptGFP signal produced represents translation of the input RNA. BHK-21 and MDBK cells are continuous cell lines known to support replication of FMDV: BHK-21 cells are used for FMDV vaccine production, whereas MDBK cells originate from a natural host of FMDV. Replication was monitored by measuring reporter expression using an IncuCyte ZOOM live cell imaging system, with analysis by our established protocols (performed 8 hours post transfection, when replication has reached its peak (Figure S2)) (Figure 2).

**Figure 2.**
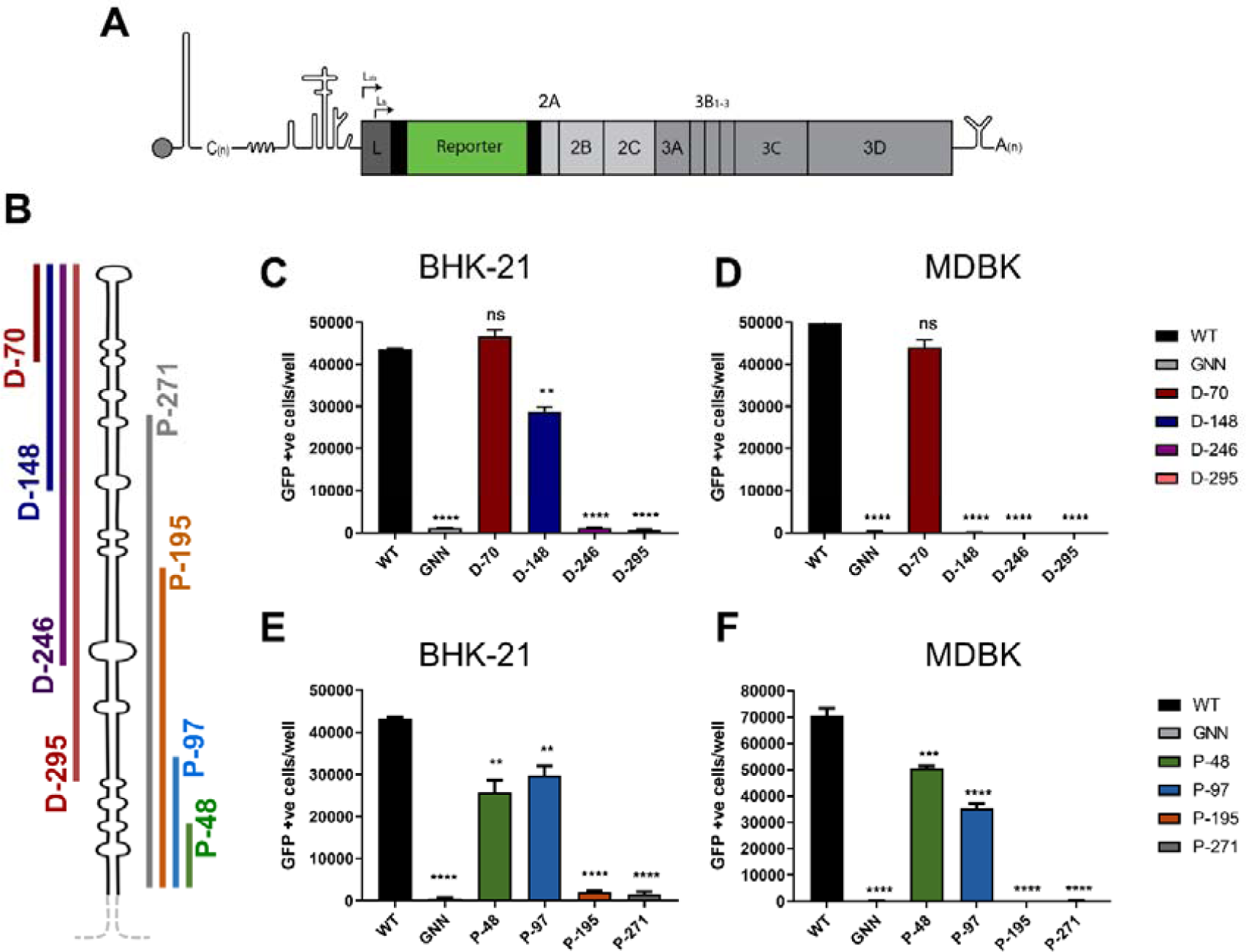
Truncations to the distal and proximal regions of the S fragment can impair replication. **(A)** Cartoon schematic of FMDV O1K replicon with the region encoding the structural proteins replaced with a GFP reporter. **(B)** Schematic representation of S fragment deletions. The maintained 3′ and 5′ proximal (P) regions are represented by the dotted grey line and was constant in all deletions. The coloured lines indicate the part of the S fragment which was removed. **(C-D)** Replication of replicons with 70, 148, 246 and 295 nucleotides removed from the distal (D) region of the S fragment was measured following transfection into BHK-21 cells or MDBK cells. Replication was monitored by GFP expression using an IncuCyte, shown at 8 hours post-transfection alongside WT and 3D^pol^ inactive mutant ‘GNN’ acting as a positive and negative control, respectively. **(E-F)** Replication of replicons with 48, 97, 195 and 271 nucleotides removed from the proximal (P) region of the S fragment and measured as in (C). (n = 3, error bars represent SEM) ** P<0.01, **** P<0.0001.

As described previously and consistent with sequence information from natural virus isolates (10, 26), the D-70 mutation caused no significant drop in replication in either cell line. However, the replicon bearing the D-148 deletion, which removes the distal half of the S fragment stem-loop, showed reduced replication in BHK-21 cells and none in MDBK cells (Figure 2C, D). Innate immunity is compromised in BHK-21 cells but intact in MDBK cells (56, 57), suggesting that the distal portion of the S fragment stem-loop plays a role in modulating a competent host immune response, in agreement with previously published data (25, 58). Replicons with the largest distal deletions, D-246 and D-295, did not replicate in either cell line.

While the consequences of deleting sequences from the distal region of the S fragment stem-loop have been reported (25), the effects of proximal deletions have not been investigated. The two smallest proximal deletions (P-48 and P-97), showed modest but significant decreases in replication compared to WT (1.7 and 1.4-fold, respectively) (Figure 2E, F). However, the larger deletions, P-195 and P-271, completely ablated replication and reporter gene expression was reduced to a level comparable to the GNN control. Interestingly, there was no significant difference between the replication of the P-48 and P-97 deletions in either BHK-21 or MDBK cells (Figure 2). Overall, the proximal part of the S fragment stem-loop appears to be more sensitive to deletion in a cell type-independent manner, with truncations as small as 48 nucleotides having a significant effect on replication.

Earlier studies indicated that the proximal part of the S fragment stem-loop is more conserved than the distal part (10). In recent years, advances in high throughput sequencing (HTS) have led to a great increase in the number of FMDV full genome sequences available on public domains, including sequences of viruses of Southern African Territories (SAT) serotypes which were previously underrepresented. This increase in the range of sequences available prompted us to revisit the subject of the S fragment variability and we carried out a covariance analysis based on 118 FMDV isolates representing all seven serotypes. Figure 3 shows that there is more base-pairing conservation in the proximal than in the distal parts of the S fragment stem-loop (Figure 3). The base-pairing conservation in the proximal part of the otherwise very variable S fragment stem-loop (low nucleotide identity of approximately 47%) suggests evolutionary pressure to maintain the structure of this portion of the stem-loop and complements our observations from the replicon experiments (Figure 2) (10, 59).

**Figure 3.**
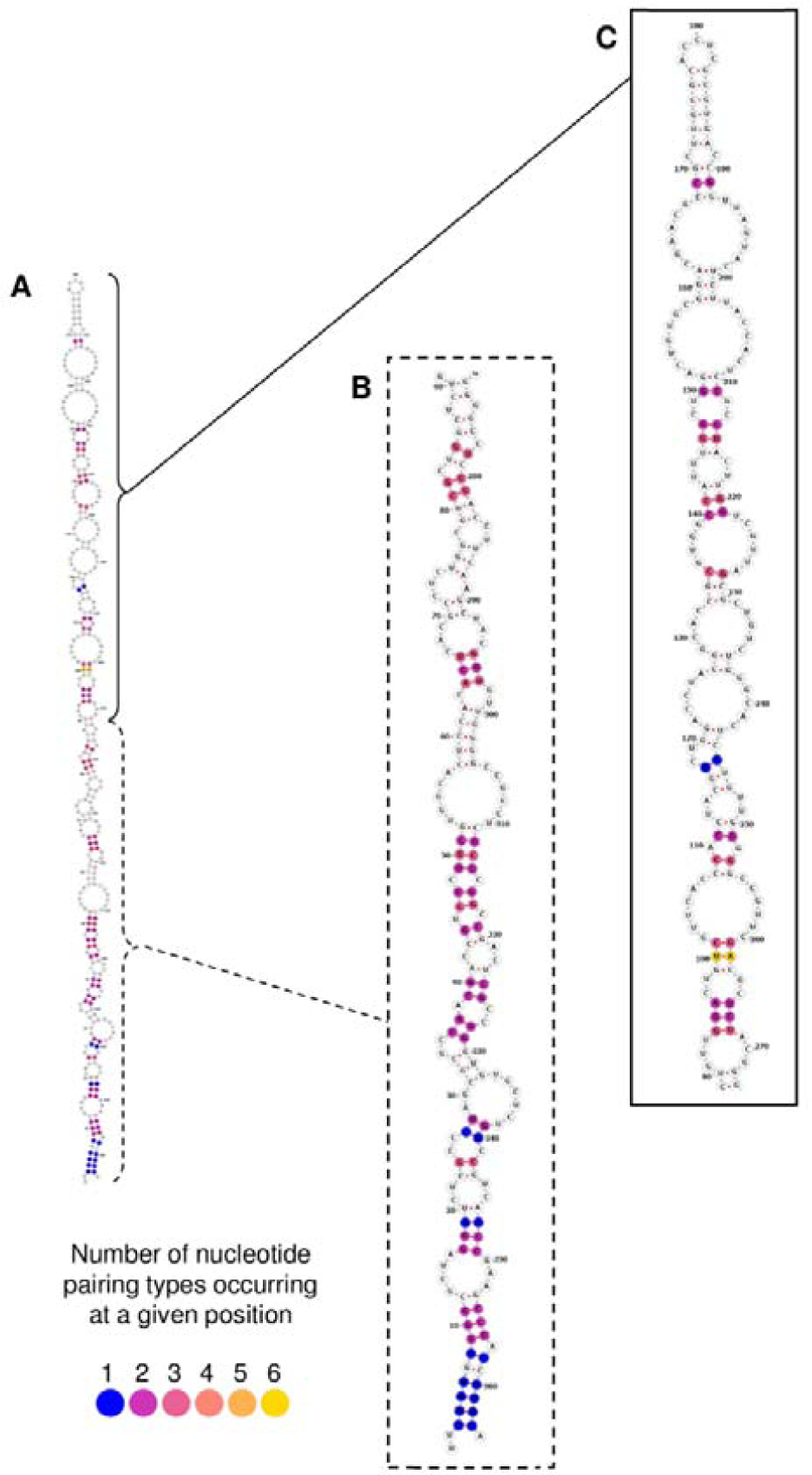
The extent of nucleotide and pairing conservation within the S fragment of the FMDV. **(A)** A schematic representation of conserved RNA structure of the S fragment in all 118 FMDV isolates. Nucleotide positions which form conserved pairing in 116/118 FMDV isolates were colour-coded according to number of pairing types (purple for one nucleotide pairing type at a given position and yellow for six nucleotide pairing types; see the colour matrix included at the left bottom corner of the figure). There are six possible nucleotide pairings: A-U, G-C, G-U, U-A, C-G and U-G. Nucleotide pairings which were not conserved in three or more FMDV isolates, remained white. Due to the length of the S fragment stem-loop, the resolution of the image was not sufficient for detailed view of the individual nucleotide pairings and so the S fragment was artificially divided into two parts, proximal and distal, and the images of their schematic RNA structures enlarged (**B-C**). **(B)** The proximal part of the S fragment included nucleotide positions 1 to 90 and 272 to 364, **(C)** while the distal part included nucleotide positions 90 - 272. Numbers represent nucleotide positions of the S fragment sequence.

### Viruses with proximal deletions of the S fragment replicate more slowly than WT and can select for a mutation in 3D^pol^ during serial passage

We investigated the consequences of proximal deletions to the S fragment stem-loop further by introducing these truncations into an FMDV O1K infectious clone. This enabled evaluation of replication in the context of the entire genome and investigation of the possible selection of compensatory or adaptive mutations during serial passage. Replicons bearing proximal deletions of the S fragment stem-loop were converted into infectious clones by replacing the ptGFP reporter sequence with the original O1K structural protein sequence. RNA transcripts of the truncated S fragment clones and WT were transfected into BHK-21 cells. Supernatants containing recovered virus were harvested and used to infect naïve BHK-21 cells for 5 continuous passages. Recovery of infectious virus from each construct was assessed by the ability of 5^th^ passage samples to induce CPE. As expected, CPE was induced and virus recovered from the WT construct. Infectious virus was also recovered from P-48 and P-97 but no virus was recovered from either P-195 or P-271 constructs.

The consequences of the P-48 and P-97 S fragment deletions for the dynamics of virus replication were investigated by assessing the time taken to induce CPE in BHK-21 cell monolayers after infection with the mutated viruses or WT at a low MOI of 0.01 PFU. The integrity of the cell monolayers was assessed at hourly intervals over 63 hours using live cell imaging monitored by IncuCyte S3 (Figure 4A). Both P-48 and P-97 viruses developed CPE more slowly than WT, as anticipated from the earlier replicon experiments (Figure 4A). For comparison, while a D-148 virus bearing a larger deletion from the distal end of the S fragment stem-loop replicated at a slower rate than the WT virus, D-148 replicated significantly faster than viruses carrying deletions at the proximal end of the S fragment (i.e., P-48 and P-97; Figure 4A); further confirming replicon experiments. Ability of the viruses to form plaques was also assessed. Both, P-48 and P-97 mutants generated significantly smaller plaques when comparing to WT and/or D-148 mutant virus (Figure 4B).

**Figure 4.**
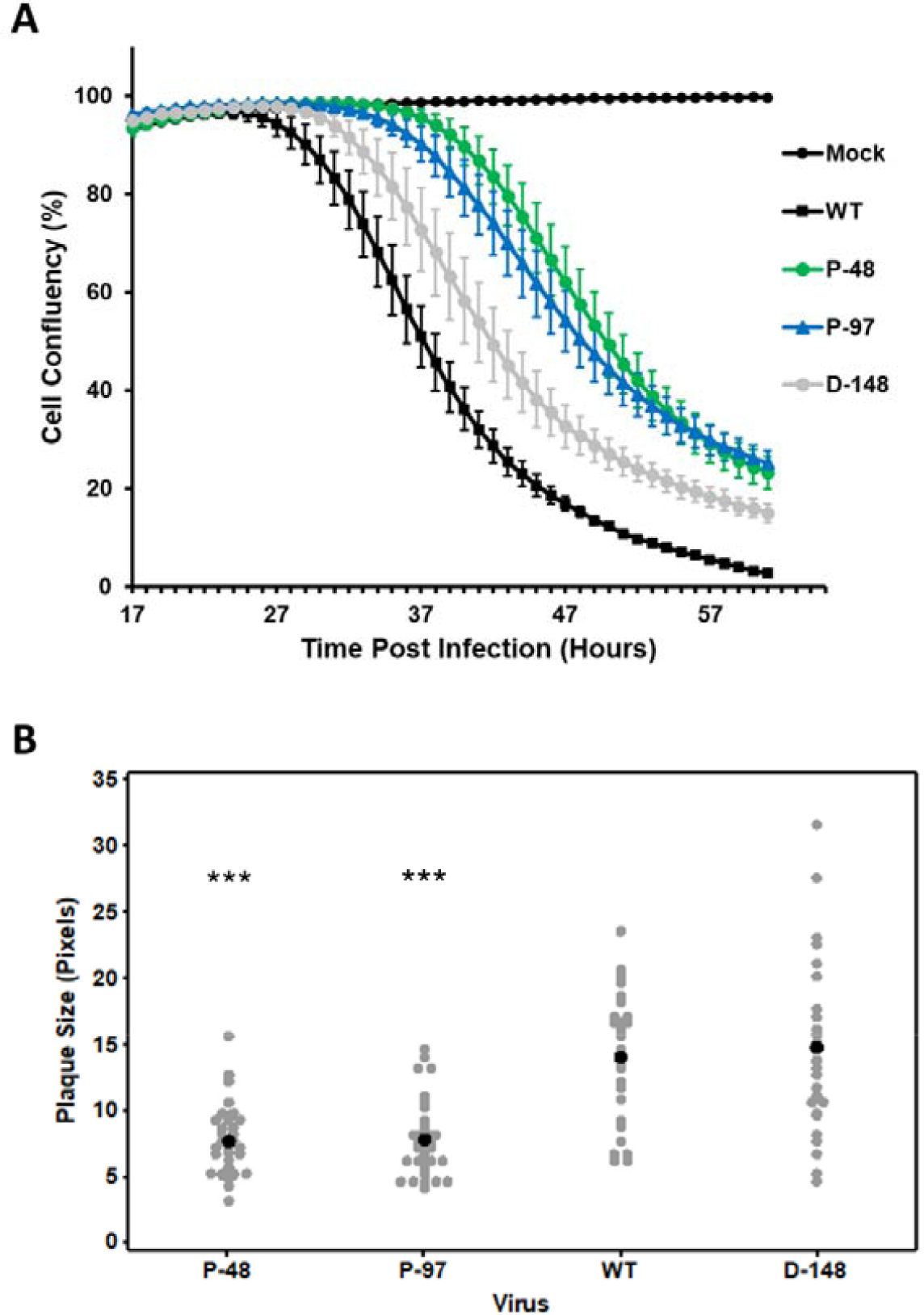
Fitness of FMDV containing a truncation at the proximal end of the S fragment. **(A)** Recovered WT, P-48, P-97 and D-148 virus populations were used to infect BHK-21 cells at an MOI of 0.01 PFU. CPE was monitored over time using an IncuCyte S3 live cell imaging and is shown as a reduction in cell confluency; (n =3 error bars represent SEM). **(B)** Plaque sizes of the WT, P-48, P-97 and D-148 variants grown on BHK-21 cells. Each well containing plaques was scanned and the plaque size was estimated in pixels. All plaques were counted to avoid bias in plaque selection. Grey dots represent individual plaques, while black dot represents the mean; *** P < 0.001 when comparing to plaque size induced by the WT.

Sequencing of recovered viruses revealed one isolate with mutations in the genomic region encoding the 2C protein, which were not investigated further. In addition, a single mutation in the 3D^pol^ sequence was found in a high proportion of the viral progeny of one of three P-97 viral replicates. The A7203C change resulted in I189L mutation in the 3D^pol^ protein amino acid sequence. The evolution of this mutation was investigated further by sequencing viruses from earlier passages. The proportion of the I189L mutation in the population increased from 40% at passage 3 to 51% at passage 4, and 61% at passage 5, suggesting it confers a selective advantage in the context of the proximal S fragment stem-loop deletion and so is sequentially enriched. With four exceptions (two isolates containing valine and two isolates containing threonine at this position) out of 1123 sequences of 3D^pol^ available, all known FMDV isolates have isoleucine at this position (Figure S3), making the amino acid substitution for leucine novel. This residue is known to interact with template RNA (Figure 5A), playing a crucial role during viral replication (60). No such mutations were observed in the WT or the P-48 mutant viruses after passage.

**Figure 5.**
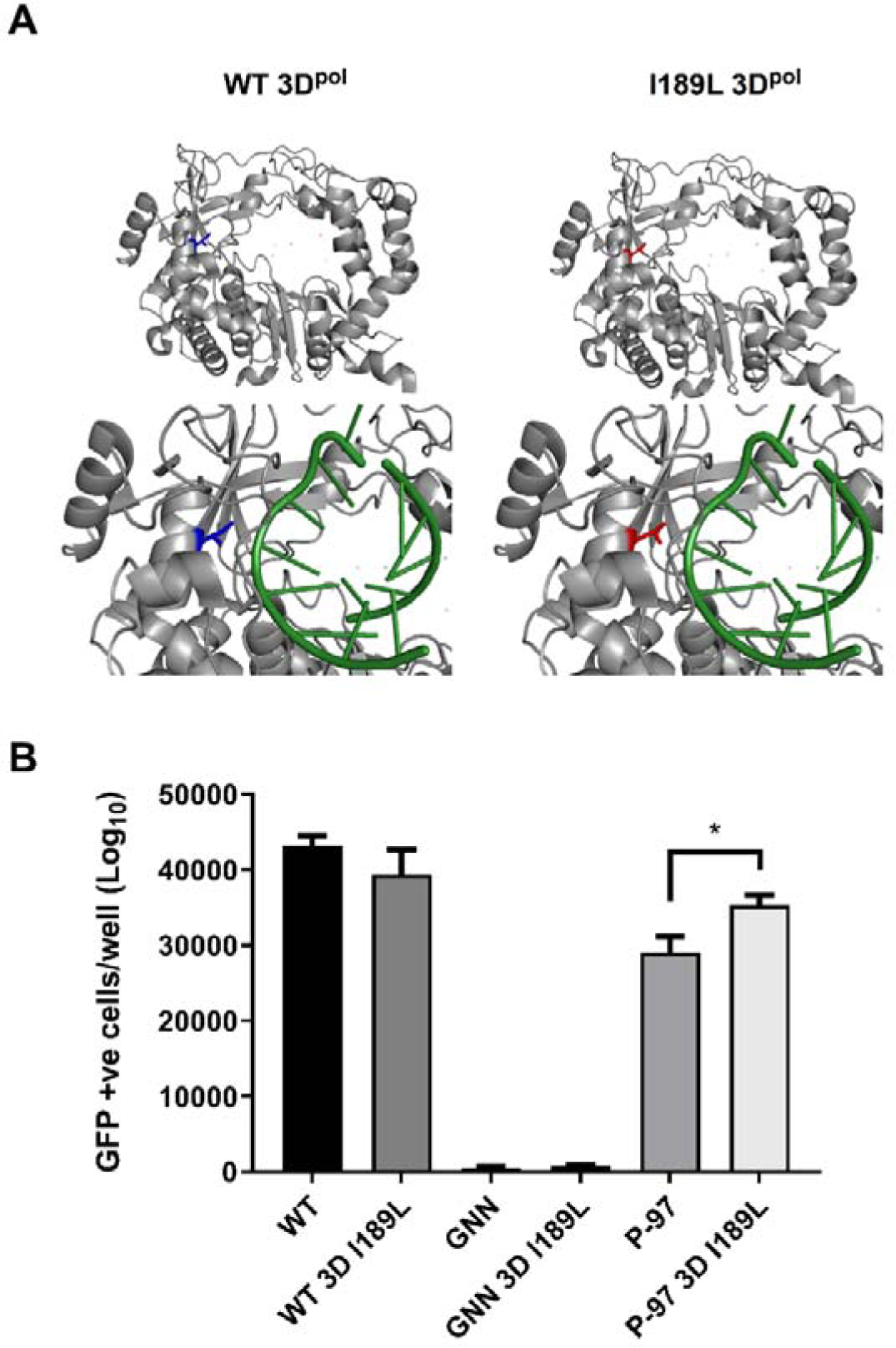
Compensatory mutation I189L in the structure of 3D^pol^ and its effect on replication of FMDV replicons. **(A)** WT 3D^pol^ crystal structure with I189 residue highlighted in blue, shown with (bottom, left) and without bound RNA (top, left). The I189L residue mutation was modelled onto the WT 3D^pol^ crystal structure and highlighted in red (right). Structure shown with (bottom, right) and without (top, right) bound RNA. PDB file 1WNE. **(B)** The 3D^pol^ I189L mutation was introduced into WT, GNN and P-97 replicons, termed WT I189L, GNN I189L and P-97 I189L respectively. Replicon RNA was transcribed and transfected into BHK-21 cells. WT and 3D^pol^ GNN replicons were included as controls, with the latter for the level of input translation (n = 3 error bars represent SEM, * P < 0.05).

### Effect of the 3D^pol^ I189L compensatory mutation on replicon replication

The 3D^pol^ I189L mutation was introduced into WT, GNN and P-97 replicons by site-specific mutagenesis and the consequence for replication assessed by reporter gene expression following transfection into BHK-21 cells, as above. Introduction of 3D^pol^ I189L into the WT replicon had no significant effect on replication, however, a small but significant increase in replication was observed when it was introduced into the P-97 replicon (Figure 5B). As expected, I189L was unable to restore the activity of the GNN negative control.

### I189L modifies 3D^pol^ activity

The influence of the 3D^pol^ I189L mutation on polymerase function was investigated using Sym/Sub assay; a highly sensitive method for interrogating polymerase activity. Sym/Sub assays have been used to study PV replication and enable extension of a radiolabelled template to be measured at the single nucleotide level (48). The method uses a 10-nucleotide template oligonucleotide which base-pairs to create a small double stranded region with a four nucleotide 5′ overhang, thus allowing 3′ nucleotide addition. In the presence of rNTPs and recombinantly expressed 3D^pol^, the addition of single or multiple nucleotides can be assessed over time by separating input template and elongation products by gel electrophoresis (Figure 6A). Initially, an assay using ATP only was undertaken to probe the ability of the enzyme to add a single nucleotide. WT and I189L 3D^pol^ proteins behaved similarly in the assay with no differences in the rate of addition of the single nucleotide (Figure 6B & C).

**Figure 6.**
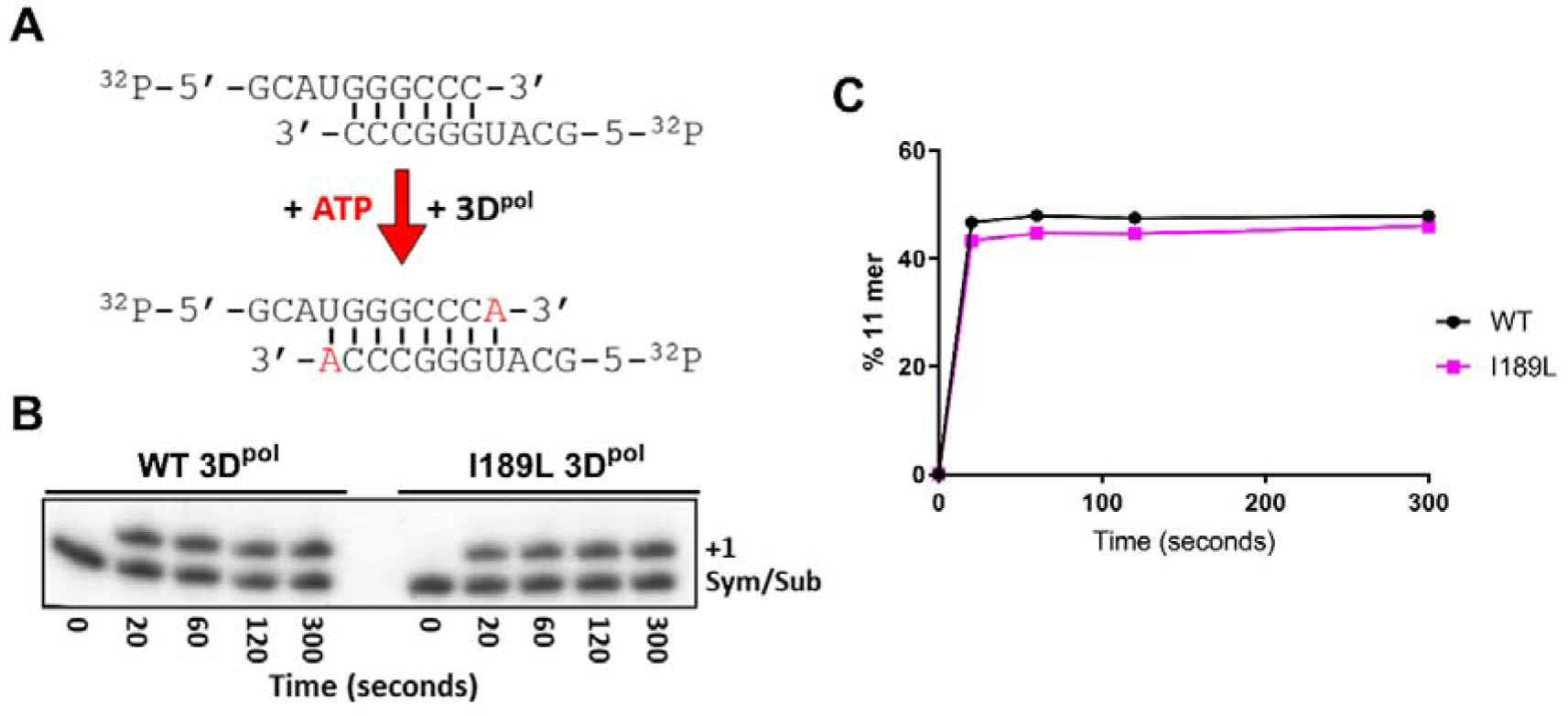
Single nucleotide addition to a Sym/Sub substrate by WT and I189L 3Dpol. **(A)** Schematic of the Sym/Sub experimental protocol. Radioactively end labelled RNA oligonucleotides are annealed before addition of rNTPs and recombinant 3D^pol^. **(B)** Extension of the 10mer Sym/Sub template with a single nucleotide (ATP) using either WT or I189L FMDV 3D^pol^. Aliquots of the reactions were taken at 20, 60, 120 and 300 seconds, RNA fragments separated by electrophoresis and visualised using a phosphoimager. **(C)** Densitometry of +1 product is plotted as rate of addition of a single nucleotide shown over time, %+1 product refers to the total amount of input Sym/Sub template that was extended by 1 nucleotide.

3D^pol^ activity was examined further by including all four rNTPs into the Sym/Sub assay. The design of the Sym/Sub oligonucleotide allows for elongation by a maximum of +4 nucleotides to produce a fully double stranded product (Figure 7A). Both the WT and the I189L 3D^pol^ enzymes produced the expected +4 product. In addition, a longer product equivalent to +12 nucleotides was produced by both polymerases (as calculated by Rf value for migration travelled) (Figure 7B). This is reminiscent of the larger products seen in recombination assays reported in the investigation of PV 3D^pol^ (48, 61). Although there was no measurable difference in rate of addition of the four nucleotides by WT or mutant enzyme, I189L 3D^pol^ produced more of the larger +12 product (Figure 7C, D). The enhanced generation of this product shows that the I189L mutation altered the activity of 3D^pol^ *in vitro* and also resulted in improved replication of the P-97 replicon (Figure 5B). These results suggest that I189L is a compensatory mutation conferring a replicative advantage to the replicon carrying the largest viable proximal truncation of the S fragment stem-loop and implies that the 3D^pol^ and the S fragment interact during replication of FMDV.

**Figure 7.**
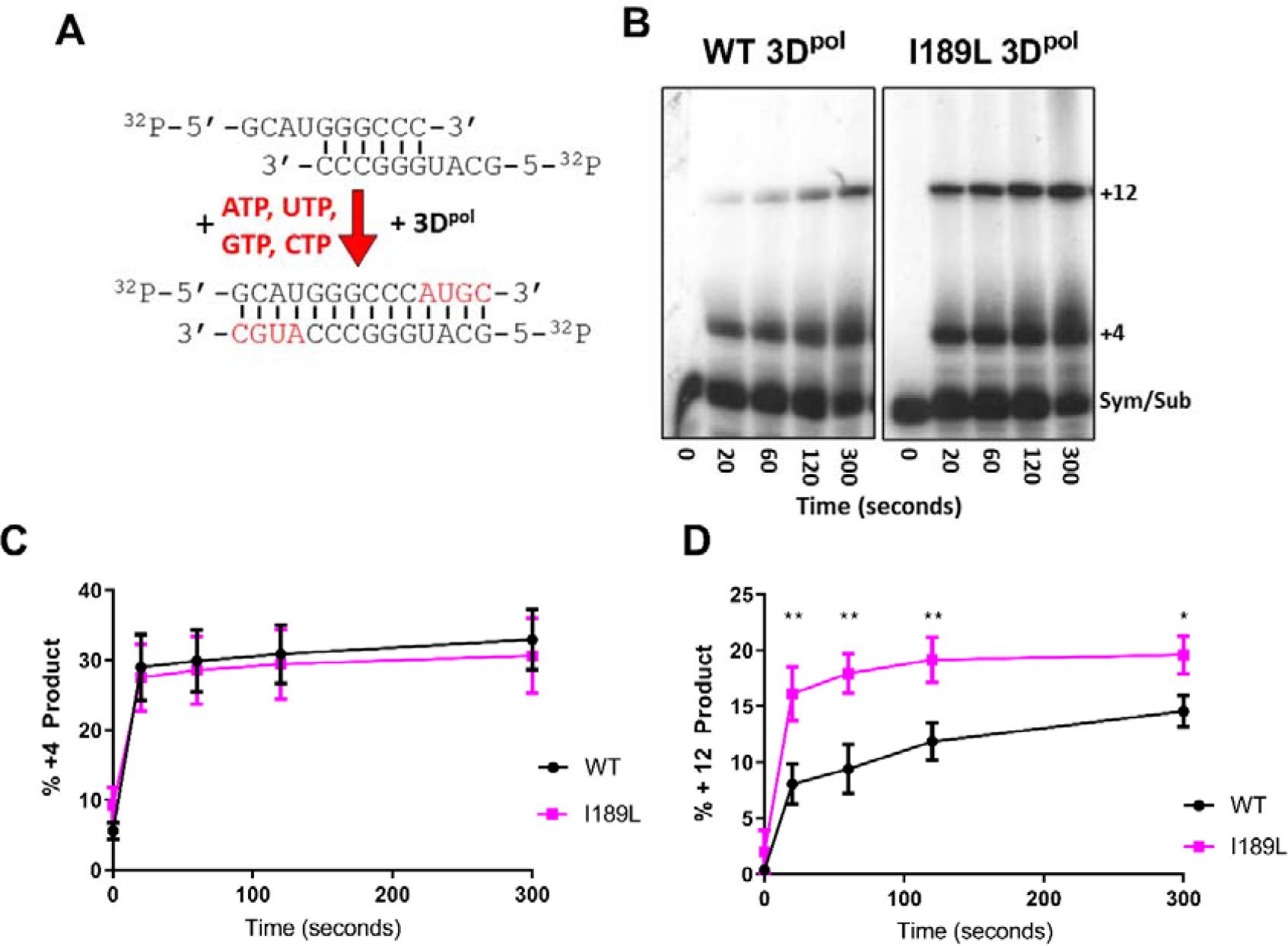
Sym/Sub assays showing differences between WT and I189L 3D^pol^ in the production of a +12 nucleotide product. **(A)** Schematic of the Sym/Sub experimental protocol using all four rNTPs. **(B)** Extension of the 10mer Sym/Sub template with all four nucleotides using either WT or I189L recombinant FMDV 3D^pol^. Aliquots of the reactions were taken at 20, 60, 120 and 300 seconds, RNA fragments separated by electrophoresis and visualised using a phosphoimager. **(C)** Densitometry of the production of the +4 product by either WT or I189L 3D^pol^ over time. (n = 4, error bars represent SEM). **(D)** Densitometry of the production of the +12 product by either WT or I189L 3D^pol^. (n = 4, error bars represent SEM). Data are shown as the % of the input Sym/Sub template elongating to the +4 or +12 products. * P < 0.05, ** P<0.01.

## Discussion

In comparison to other picornaviruses, the 5′ UTR of FMDV is uniquely large and complex, comprising several distinct structural domains, the functions of most of which are poorly understood. The first *c.* 360 nucleotides are predicted to form a single, long stem-loop called the S fragment (22). Although supported by functional studies, to our knowledge the secondary structure of the S fragment has not been determined biochemically (25). Here, using SHAPE chemistry applied to a sequence of a well-studied FMDV isolate (O1 Kaufbeuren) (35), we confirm that the S fragment folds into a single, long stem-loop containing several bulges formed by unpaired nucleotides. The importance of some of these bulges for viral viability was determined previously (25).

Despite the high sequence and length variability of the S fragment observed in FMDV field isolates, most viruses maintain the full-length stem-loop (10). This suggests a strong evolutionary pressure to maintain the structure. The S fragment is known to interact with viral and cellular proteins and the 3′ UTR, likely facilitating circulation of the viral genome during replication (64–66). However, there is evidence that some deletions to the distal part of the S fragment are tolerated and a number of unrelated field isolates from different FMDV serotypes (e.g., A, C and O) have been shown to carry deletions in their S fragments. These deletions arose independently and range in length from few to 76 nucleotides (10, 26, 62). Unlike deletions found in other parts of the FMDV genome (e.g. deletions found in the region encoding 3A), those found in the S fragment do not appear to be host specific, with viruses carrying deletions within the S fragment being isolated from both cattle and pigs (10, 26, 62, 63). A more recent study showed that deletion of 164 nucleotides from the distal part of the S fragment can produce a viable, although attenuated virus, suggesting a role in evasion of innate immunity, at least partially explaining the evolutionary trend to maintain the full structure of the S fragment by field viruses (25). Here, we show that although deletion of 246 nucleotides or more from the distal region on the S fragment was not tolerated, mutants carrying deletions D-70 and D-148 remained replication-competent.

The proximal part of the S fragment stem-loop shows higher sequence and nucleotide pairing conservation compared to the distal region (Figure 3). Surprisingly, deletions to the proximal part of the stem-loop (i.e., up to 97 nucleotides) were also tolerated, although to a lesser extent than distal deletions, resulting in both attenuated replicons and viruses (cf P-48 and P-97 mutants versus D-148 mutant, Figure 2 and 4) (10). Interestingly, while the replication of D-148 mutant replicon was more impaired in MDBK cells than in the BHK-21 cells, this cell-specific difference was not seen for the viable replicons carrying deletions to the proximal part of the S fragment stem-loop. Since, unlike BHK-21 cells, MDBK cells appear to have a functional interferon pathway this might suggest that, although required for viral replication, this proximal part of the S fragment stem-loop does not play a part in the evasion of the innate immunity (56, 57).

Of three viral isolates carrying the maximum viable proximal deletion to the S fragment (P-97), two resulted in selective enrichment of a compensatory mutation over serial passages. In both cases, the mutations showed a trend towards fixation. One isolate developed a number of mutations in 2C (data not shown), while another developed a single I189L mutation in a highly conserved site of 3D^pol^ known to form hydrophobic contact with the sugar-phosphate backbone of the template RNA during viral replication (60). Given the essential role of I189 in viral RNA replication, we investigated the effect of the I189L mutation on replication and RNA strand synthesis. Introduction of I189L into the P-97 replicon enhanced replication, suggesting an advantageous effect of this mutation. Biochemical analysis of the I189L 3D^pol^ mutant did not indicate an altered rate of nucleotide incorporation when compared to the WT polymerase. However, the synthesis of a larger than predicted +12 product was greater with the I198L 3D^pol^ enzyme compared to WT. This suggests that the I198L mutation may influence the fidelity of the polymerase, possibly due to an altered interaction between the amino acid side chain and the RNA template. Extended products have been observed in similar reactions with the 3D^pol^ of PV and were proposed to be the result of a nucleotide misincorporation, facilitating a template switch (48, 67, 68). In addition, it has been reported that greater fidelity of viral RNA polymerases is associated with increased insertion/deletion (indel) events during viral replication (69). Whether the longer product observed here occurred as the result of strand slippage, or a template switch remains to be investigated. Nevertheless, slow accumulation of the compensatory mutation over serial passage in a virus carrying maximal viable proximal deletion to the S fragment structure, and the small advantageous effect of this mutation on replicon replication suggests the possibility of (direct or indirect) interaction between the S fragment and the viral polymerase (or its precursor). Further studies are required to verify this interplay.

In conclusion, using biochemical methods, we confirmed a previously proposed secondary structure of the S fragment of FMDV. Despite its higher sequence and nucleotide pairing conservation, small deletions to the proximal part of the S fragment stem-loop are viable, although resulting in attenuated replicons and viruses. An advantageous compensatory mutation within a highly conserved site of 3D^pol^ appeared during serial passage of a viral isolate bearing the largest viable mutation to the proximal part of the S fragment, suggesting potential interactions between the S fragment and the viral polymerase during replication.

## Data availability

Additional information and data underpinning the work described here (e.g. primer sequences) can be obtained by contacting the corresponding authors.

