## Supplementary data for "The dual role of a highly structured RNA (the S fragment) in the replication of foot-and-mouth disease virus"

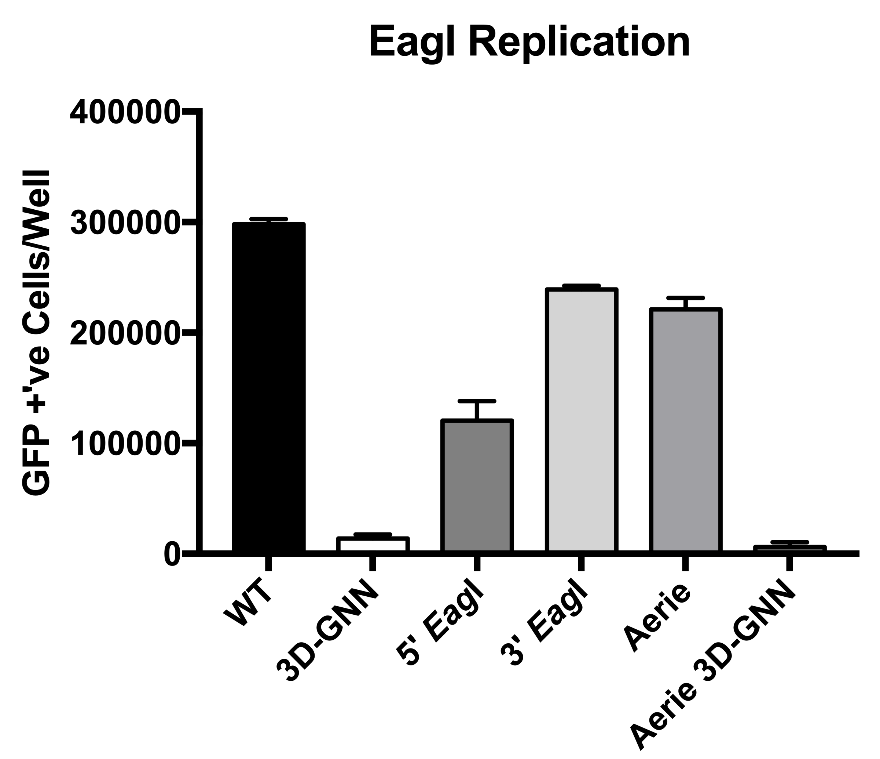

**Supplementary Figure S1. Effect of introduction of *Eag*I sites on replication.** Number of GFP positive cells/well detected at 8 hours post-transfection of BHK-21 cells. WT represents the original O1K replicon sequence. 3D-GNN is the polymerase defective negative control. 5′ *Eag*I replicon possesses mutations to the sequence to create an *Eag*I site at the 5′ end of the S fragment. 3′ *Eag*I possesses mutations to the sequence to create and *Eag*I site at the 3′ end of the S fragment. Aerie shows replicon with both 5′ and 3′ sites modified and Aerie 3D-GNN comprises both *Eag*I sites and 3D-GNN mutation. n=2, error bars represent SEM.

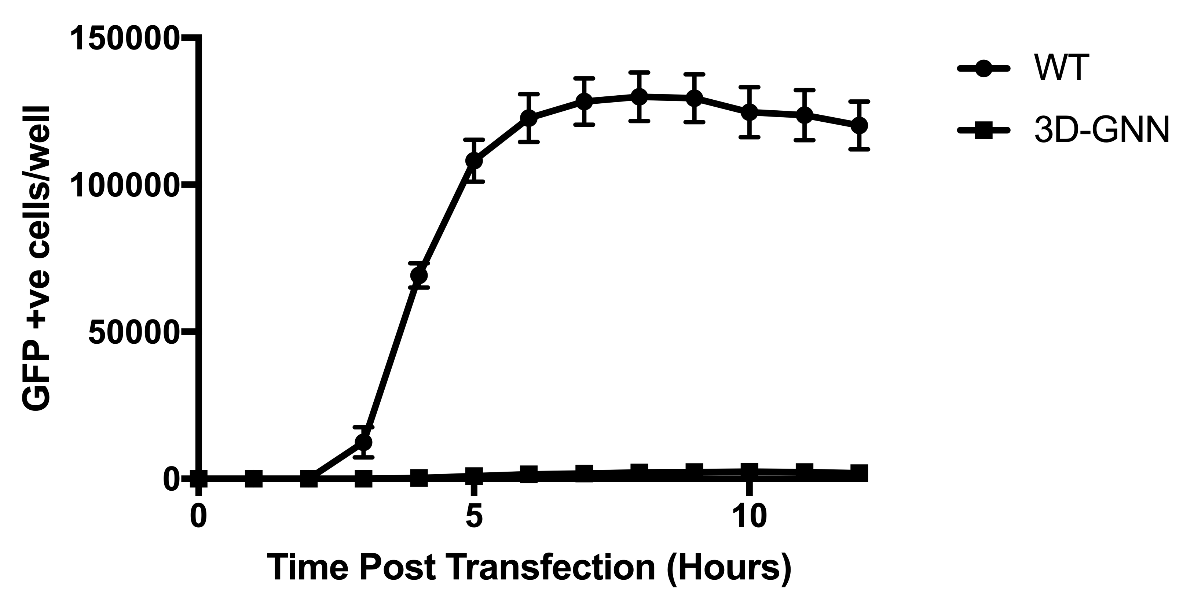

**Supplementary Figure S2. Time course of FMDV replicon replication.** Number of GFP positive cells/well detected at hourly time points post-transfection of WT and 3D-GNN replicons in BHK-21 cells. n=2, error bars represent SEM.

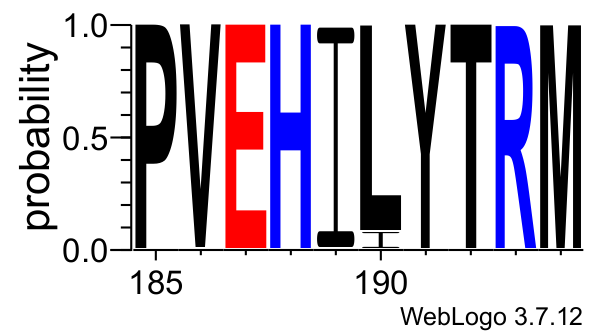

**Supplementary Figure S3. Conservation of the isoleucine at the position 189 (I189) within the 3D^pol^ protein of FMDV.** Sequence logo based on 1123 sequences of FMDV 3D^pol^ available on fmdbase.org and prepared using WebLogo 3 server. The x-axis represents amino acids positions of the 3D^pol^ protein of FMDV, while the y-axis shows probability of a particular amino acid being present at a given position. Amino acids are colour-coded according to their charge.

**Supplementary Table S1**

FMDV sequences selected from GenBank

| **Serotype:** | **Number of isolates:** | **GenBank accession numbers:** |
| --- | --- | --- |
| A | 19 | AY593788, MH053305, JF749843, HM854024, HQ832580, MH053306, KM268896, AY593802, KJ608371, MH053307, AY593751, AY593754, AY593761, AY593764, AY593766, AY593767, HM854022, AY593791, AY593794 |
| Asia 1 | 12 | AY593795, AY687334, DQ533483, DQ989306, DQ989315, DQ989319, EF149010, EF614458, HQ632774, JF739177, KM268898, MF782478 |
| C | 6 | MH053308, KM268897, MH053309, AJ133357, MH053310, AJ007347 |
| O | 21 | AY593819, MH053313, MH053311, MH053312, KF112885, KJ206909, HQ632769, HQ632771, KU291242, KR401154, GU384683, KF694737, AJ539140, MH053315, JX040491, MH053317, MH053318, MH053316, KJ560291, DQ404170, KU821591 |
| SAT 1 | 21 | AY593838, AY593845, MH053319, AY593844, JF749860, MH053321, AY593846, AY593839, AY593842, AY593841, AY593840, MH053322, AY593843, KM268899, MH053323, MH053324, MH053325, MH053326, MH053327, MW355668, MW355669 |
| SAT 2 | 19 | MH053330, MH053332, MH053328, MH053329, JX014255, MH053333, AY593849, JX014256, MW355670 - MW355673, AY593847, MH053335, KM268900, JF749862, MH053336, MH053337, KU821592, |
| SAT 3 | 20 | AY593853, AY593851, MH053339, MH053340, MH053344, MH053343, AY593850, KJ820999, MH053341, MH053351, KX375417, KM268901, MH053350, MW355674 - MW355680 |
